# Genuine beta bursts in human working memory: controlling for the influence of lower-frequency rhythms

**DOI:** 10.1101/2023.05.26.542448

**Authors:** Julio Rodriguez-Larios, Saskia Haegens

## Abstract

Human working memory is associated with significant modulations in oscillatory brain activity. However, the functional role of brain rhythms at different frequencies is still debated. Modulations in the beta frequency range (15–40 Hz) are especially difficult to interpret because they could be artifactually produced by (more prominent) oscillations in lower frequencies that show non-sinusoidal properties. In this study, we investigate beta oscillations during working memory while controlling for the possible influence of lower frequency rhythms. We collected electroencephalography (EEG) data in 31 participants who performed a spatial working-memory task with two levels of cognitive load. In order to rule out the possibility that observed beta activity was affected by non-sinusoidalities of lower frequency rhythms, we developed an algorithm that detects transient beta oscillations that do not coincide with more prominent lower frequency rhythms in time and space. Using this algorithm, we show that the amplitude and duration of beta bursts decrease with memory load and during memory manipulation, while their peak frequency and rate increase. In addition, interindividual differences in performance were significantly associated with beta burst rates. Together, our results show that beta rhythms are functionally modulated during working memory and that these changes cannot be attributed to lower frequency rhythms with non-sinusoidal properties.

## Introduction

Working memory refers to the capacity of holding and manipulating information, which is fundamental for any type of goal-directed behavior (Baddeley, 2010; G. A. Miller et al., 1960). In order to study the behavioral and neural correlates of working memory in humans, a wide variety of working-memory tasks have been developed (Repovš & Baddeley, 2006). Working-memory tasks usually involve a delay or maintenance period in which information (typically in a specific sensory modality) is transiently kept in mind in the absence of external stimulation. Some working-memory tasks also allow studying the effect of memory load (i.e., number of items to be remembered) and memory manipulation (i.e., the modification of memory items in addition to their maintenance). The combination of these tasks with neuroimaging and electrophysiological techniques has allowed the identification of distinct brain mechanisms supporting working memory (D’Esposito & Postle, 2015; E. K. Miller et al., 2018).

Previous research in humans has shown that working memory is associated with significant modulations in oscillatory activity (Pavlov & Kotchoubey, 2020b). Oscillatory activity, as measured with the Electroencephalogram (EEG), originates from the summed activity of pyramidal neurons arranged perpendicular to the scalp (Cohen, 2017), and is thought to reflect local excitability and long-range communication (Klimesch et al., 2007). Previous work on the role of brain oscillations in working memory has mostly focused on frontal theta (∼4–8 Hz) and posterior alpha (∼8–14 Hz) oscillations, which form the most prominent rhythms in the human EEG (Klimesch, 1999). Modulations in the beta range have received considerably less attention, with the few studies reporting beta power modulations highly inconsistent regarding the direction and topography of such effects (Pavlov & Kotchoubey, 2020b). In addition, while changes in alpha and theta center frequency have shown to be functionally relevant (Angelakis et al., 2004; Rodriguez-Larios & Alaerts, 2019), frequency modulations in the beta range have not yet been investigated in the context of human working memory.

The detection of beta oscillations in EEG/MEG signals is not trivial. Relative to oscillations in lower frequencies, beta oscillations are less sustained (Sherman et al., 2016) and have lower amplitudes (Pfurtscheller & Cooper, 1975). Moreover, changes in the beta range can be artifactually produced by (more prominent) lower frequency oscillations with non-sinusoidal properties (Schaworonkow, 2023; Schaworonkow & Nikulin, 2019). Non-sinusoidal rhythms show a peak in the frequency spectrum at their main (actual) frequency and additional peaks at frequencies approximating their harmonics. Two well-known examples are the somatosensory *mu* rhythm (with a main peak around 10 Hz and a second harmonic around 20 Hz; (Kulhman, 1978) and the frontal sawtooth theta rhythm (with a main peak around 6 Hz and a third harmonic around 18 Hz; Onton, Delorme, & Makeig, 2005). Consequently, if beta rhythms co-occur in time and space with (more prominent) lower frequency rhythms, we cannot rule out the possibility that they are artifactually caused by non-sinusoidalities of the lower frequency rhythm. Since this is not typically controlled for in spectral analysis, previous literature on the role of beta in working memory is hard to interpret. In fact, the most recent review on the EEG/MEG correlates of working memory concludes that (at least part of) the reported beta modulations are likely to reflect an artifactual harmonic of lower frequency rhythms (Pavlov & Kotchoubey, 2020b).

In this study, we assessed whether beta oscillations are significantly modulated during working memory. For that purpose, we recorded 96-electrode EEG while participants (N=31) performed a working-memory task in which they had to remember and manipulate the spatial location of visual stimuli. In order to rule out the possibility that modulations in the beta range reflect non-sinusoidal rhythms in lower frequencies, we developed an algorithm that detects beta oscillatory events that do not co-occur in time and space with more prominent oscillatory events in lower frequencies. We extracted four beta burst parameters (amplitude, duration, frequency and rate) and assessed their modulation in relation to memory retention, cognitive load, memory manipulation and behavioral performance.

## Materials and Methods

### Participants

31 healthy adult participants (13 male) took part in the experiment. The mean age was 32.5 years (SD = 8.5). Participants reported normal or corrected-to-normal vision and no history of neurologic or psychiatric diagnosis. Informed consent procedure and study design were approved by the Institutional Review Board (IRB) of the New York State Psychiatric Institute. Participants were compensated for their participation (at 25 USD per hour).

### Design and task

Participants performed a visual working-memory task while EEG was recorded (see **Figure 1A**). First a fixation cross was presented for 1 s. Then, participants were presented with one (or three) angles in a circle. Following a delay period of 3 s, a cue was presented for 1 s. The cue was either ‘stay’, meaning that the correct answer was the presented angle, or ‘switch’, indicating that the correct answer was the opposite angle. After the instruction, a response mapping diagram was shown, indicating which button number (1 to 8) corresponded to which angle. This response map was randomized in each trial. After each response, participants received feedback based on their answer (a green circle for correct answers and a red circle for incorrect answers). Participants performed four blocks of 48 trials in approximately 1 hour (192 trials per participant).

**Figure 1.**
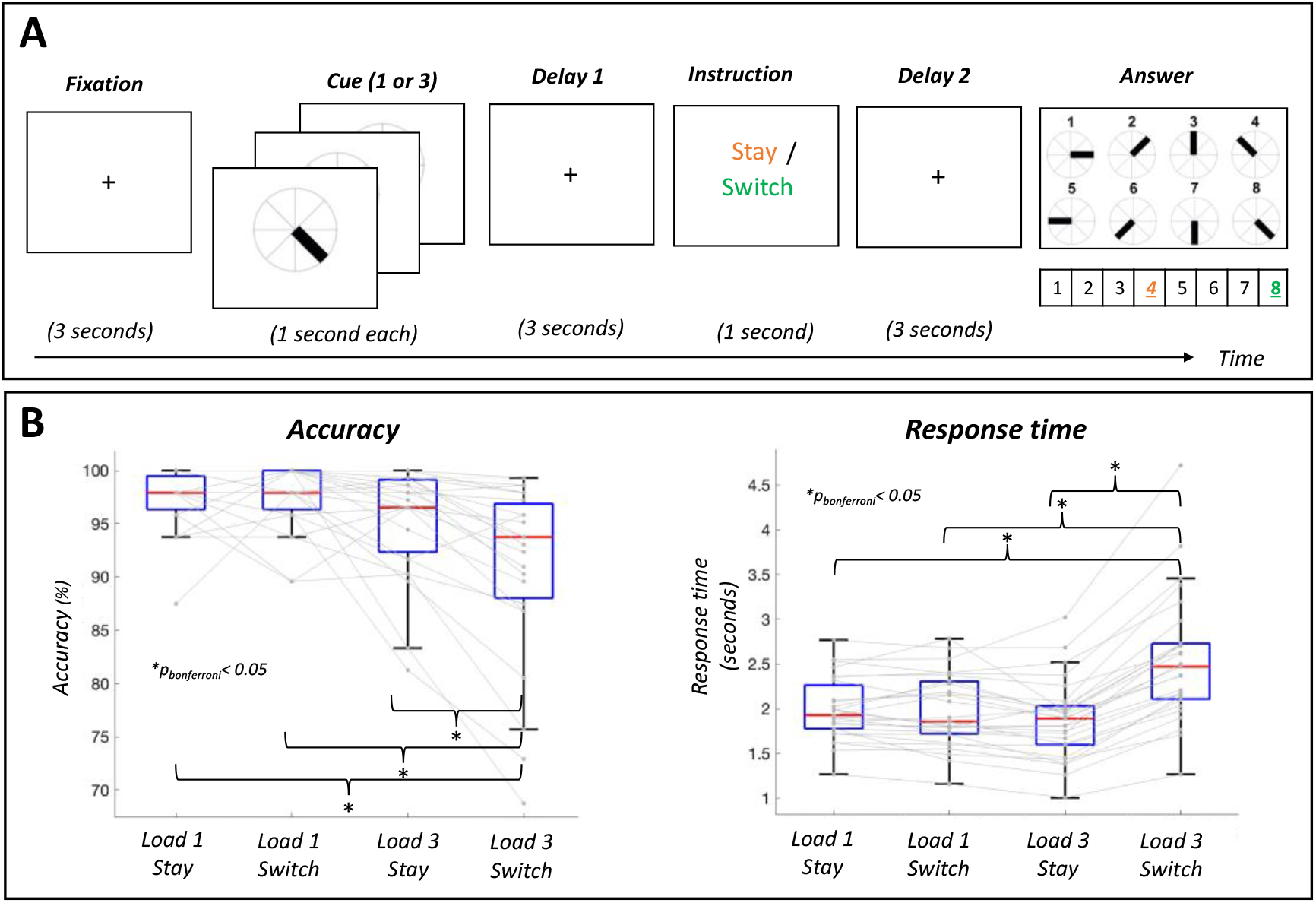
Spatial working memory task and behavioral performance. **A)** Participants were asked to remember the angle of one or three visual stimuli (*Load 1* or *Load 3*). Based on the subsequent instruction, they had to report the presented angle (*Stay*) or its opposite (*Switch*) using a computer keyboard. **B)** Participants showed significantly lower accuracy and higher response times in the condition *Load 3 Switch* relative to the other three conditions (*Load 1 Stay, Load 1 Switch, Load 3 Stay*).

### EEG acquisition

96-electrode scalp EEG was collected using the BrainVision actiCAP system (Brain Products GmbH, Munich, Germany) with a sampling rate of 500 Hz. Electrodes were labeled according to the international 10-20 system and the reference electrode during the recording was Cz. Amplification and digitalization of the EEG signal was done through an actiCHamp DC amplifier (Brain Products GmbH, Munich, Germany) linked to BrainVision Recorder software (version 2.1, Brain Products GmbH, Munich, Germany). Vertical (VEOG) and horizontal (HEOG) eye movements were recorded by placing additional bipolar electrodes above and below the left eye (VEOG) and next to the left and right eye (HEOG). In addition, electrocardiogram (ECG) was also recorded using bipolar electrodes.

### EEG preprocessing

Pre-processing was performed in MATLAB R2021a using custom scripts and functions from EEGLAB (Delorme & Makeig, 2004) and Fieldtrip (Oostenveld et al., 2011) toolboxes. Data were first resampled to 250 Hz and filtered between 1 and 40 Hz. Noisy electrodes were automatically detected (EEGLAB function *clean_channels*) and interpolated. A mean of 9.7 channels (SD = 6.5) were interpolated. EEG data were re-referenced to the common average and independent component analysis (*runica* algorithm) was performed. An automatic component rejection algorithm (IClabel) was employed to discard components associated with muscle activity, eye movements, heart activity or channel noise (threshold = 0.8; see Pion-Tonachini et al., 2019). In addition, components with an absolute correlation with HEOG, VEOG or ECG channels higher than 0.8 were discarded. The mean number of rejected components was 18.1 (SD = 5.7). Furthermore, Artifact Subspace Reconstruction (ASR) was employed to correct for abrupt noise with a cutoff value of 20 SD (Chang et al., 2020). Four participants were excluded due to uncorrectable EEG artifacts or technical problems during data acquisition.

### Beta burst detection algorithm

Trial EEG data were transformed to the time-frequency domain using 6-cycles Morlet wavelets (as implemented in the matlab function BOSC_tf; see Whitten et al., 2011). Amplitude at each frequency (1 to 40 Hz) was extracted from the real component of the convolution between the EEG signal and the family of wavelets. We used an estimate of 1/f aperiodic activity as amplitude threshold to detect oscillatory bursts (**Figure 2A**). Aperiodic activity was estimated per electrode and participant by fitting a straight line in log–log space to the average EEG frequency spectrum after excluding frequencies forming the maximum peak (for similar approaches, see Caplan et al., 2015; Goyal et al., 2020; Kosciessa et al., 2020). Oscillatory bursts were defined as time points in which the amplitude at a specific frequency exceeded the estimate of aperiodic activity for at least a cycle. In order to rule out the possibility that the detected oscillatory bursts artifactually originated from broadband changes or from a different rhythm at another frequency, only oscillatory bursts that formed the peak with the greatest prominence of the 1/f-subtracted frequency spectrum were selected (see **Figure 2B**). Using this algorithm, four burst parameters were estimated for the beta range: i) amplitude: mean prominence of the peaks of the detected bursts after subtracting aperiodic activity, ii) duration: mean number of cycles of detected oscillatory bursts, iii) rate: mean number of oscillatory bursts, and iv) frequency: mean peak frequency of detected oscillatory bursts (see **Figure 2C** for topography of each of the extracted parameters when averaging across conditions and participants). MATLAB code of the beta burst algorithm can be accessed here: https://osf.io/fkhnb/?view_only=13d1c0ac76574be6a1ba7f300fdd98e4

**Figure 2.**
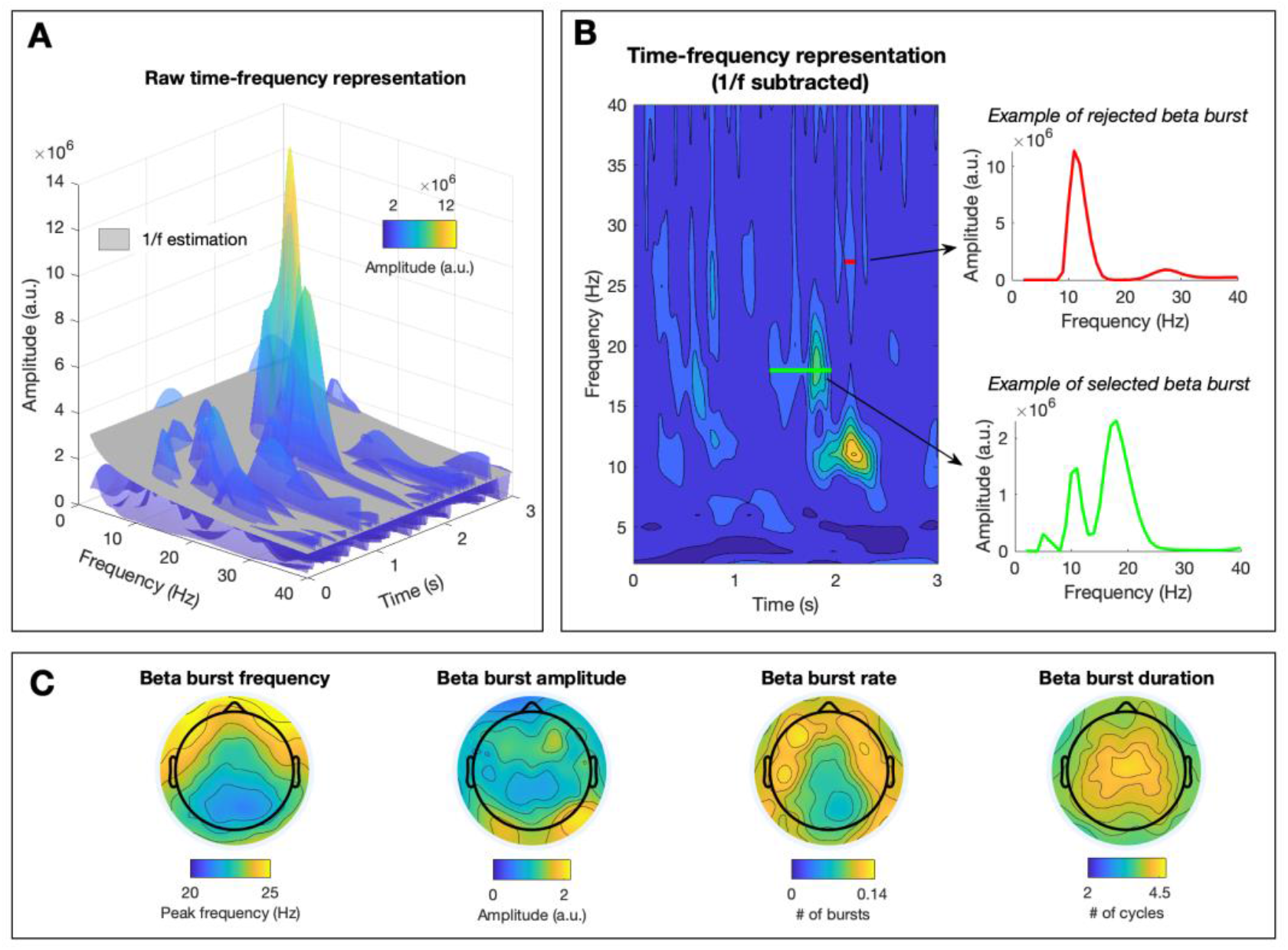
Beta burst detection algorithm and topography of beta burst parameters. **A)** Time-frequency representation of a trial. The estimate of aperiodic activity (in grey) was used as power threshold to detect beta bursts. **B)** Depiction of the time-frequency representation of the same trial after subtraction of the estimate of 1/f aperiodic activity (left panel), and frequency spectra of two exemplary beta bursts (right panels). In order to rule out the possibility that detected beta bursts are caused by the non-sinusoidal properties of lower frequency rhythms, only bursts that show a maximum spectral peak in the beta range were selected. An example of a selected burst is depicted in green and an example of a rejected burst is depicted in red. **C)** Topographical representation of average values of each of the extracted burst parameters (frequency, amplitude, rate and duration) across participants and conditions.

### Statistical analysis

For the behavioral analysis, a two-way repeated-measures ANOVA and post-hoc t-tests were performed with the JASP software (Love et al., 2019). Cognitive load (3 items vs 1 item) and memory manipulation (switch vs stay instructions) were defined as factors. This analysis was performed for accuracy and reaction times separately. The effect size was estimated with eta squared (η^2^) and Bonferroni correction was applied for multiple comparisons.

For the EEG data, a cluster-based permutation statistical test (Maris & Oostenveld, 2007) was used to assess condition-related differences in each beta burst parameter. This test controls for the type I error rate arising from multiple comparisons using a non-parametric Montecarlo randomization and taking into account the dependency of the data. First, cluster-level test statistics are estimated in the original data and in 1,000 permutations of the data. Cluster-level test-statistics are defined as the sum of t-values with the same sign across adjacent electrodes that are above a specified threshold (i.e., 97.5th quantile of a T-distribution). Then, the cluster-level statistics from the original data were evaluated using the reference distribution obtained by taking the maximum cluster t-value of each permutation. Cluster-corrected *p-values* are defined as the proportion of random partitions whose cluster-level test statistic exceeded the one obtained in the observed data. Significance level for the cluster permutation test was set to 0.025 (corresponding to a false alarm rate of 0.05 in a two-sided test). Paired-samples t-test was chosen as the first-level statistic to compare experimental conditions (*Delay* vs *Fixation, Load 3* vs *Load 1* and *Switch* vs *Stay*), and the *Pearson* correlation coefficient was chosen to assess correlations between each of the beta bursts parameters and performance (reaction time and accuracy). The effect size was estimated for each significant cluster with Cohen’s *d*.

## Results

### Behavioral performance

Repeated measures ANOVA on accuracy revealed a significant main effect of cognitive load (*F*(1,26) = 14.05 ; *p* < .001; η^2^= 0.23), a significant main effect of memory manipulation (*F*(1,26) = 11.34 ; *p* < 0.001; η^2^= 0.048) and a significant cognitive load by memory manipulation interaction (*F*(1,26) = 16.44; *p* < .001; η^2^= 0.070). Post-hoc t-tests showed that cognitive load only affected accuracy in the manipulation condition (lower accuracy in *Load 3 Switch* relative to *Load 1 Switch*) (*t*(26) = 5.17; ; *p*_*bonf*_ < .001) while memory manipulation only affected accuracy when load was high (lower accuracy in *Load 3 Switch* relative to *Load 3 Stay*) (*t*(26) = 5.24; *p*_*bonf*_ < .001) (see **Figure 1B** left panel). In addition, participants showed significantly less accuracy in high load with manipulation condition (*Load 3 Switch*) relative to the low load without manipulation condition (*Load 1 Stay*) (*t*(26) = 4.86 ; *p*_*bonf*_ < .001).

Similarly, repeated measures ANOVA on reaction time revealed a significant main effect of cognitive load (*F*(1,26) = 13.15; *p* = .001; η^2^= 0.11), a significant main effect of memory manipulation (*F*(1,26) = 79.27 ; *p* < .001; η^2^= 0.24) and a significant cognitive load by memory manipulation interaction (*F*(1,26) = 98.82; *p* < .001; η^2^= 0.26). Post-hoc t-tests showed that cognitive load only affected response times in the manipulation condition (lower reaction time in *Load 3 Switch* relative to *Load 1 Switch*) (*t*(26) = -8.02; ; *p*_*bonf*_ < .001) while memory manipulation only affected response times when load was high (lower reaction time in *Load 3 Switch* relative to *Load 3 Stay*) (*t*(26) = -13.29; *p*_*bonf*_ < .001) (see **Figure 1B** right panel). Participants also showed significantly higher response times in high load with manipulation condition (*Load 3 Switch*) relative to the low load without manipulation condition (*Load 1 Stay)* (*t*(26) = -7.66; *p*_*bonf*_ < .001).

### Beta burst modulations with memory retention

Contrasting the first delay window with the fixation window, we assessed beta burst modulations associated with memory retention and found a significant decrease in beta burst amplitude (*t*_*cluster*_ = -341.30; *p*_*cluster*_ < .001; *d* = 0.40; **Figure 3A**), a significant decrease in beta burst duration (*t*_*cluster*_ = -35.82; *p*_*cluster*_ = .002; *d* = 0.40; **Figure 3B**) and significant increase in beta burst frequency (*t*_*cluster*_ = 87.56; *p*_*cluster*_ = .002; *d* = 0.33; **Figure 3C**). The topographical distribution of significant modulations was widespread for burst amplitude and burst frequency. On the other hand, the effect on burst duration was limited to centro-parietal electrodes. No significant differences were found for beta burst rate (*p*_*cluster*_ > .05; **Figure 3D**). The average frequency spectra of beta bursts in each condition across electrodes are depicted in **Figure 3E**.

**Figure 3.**
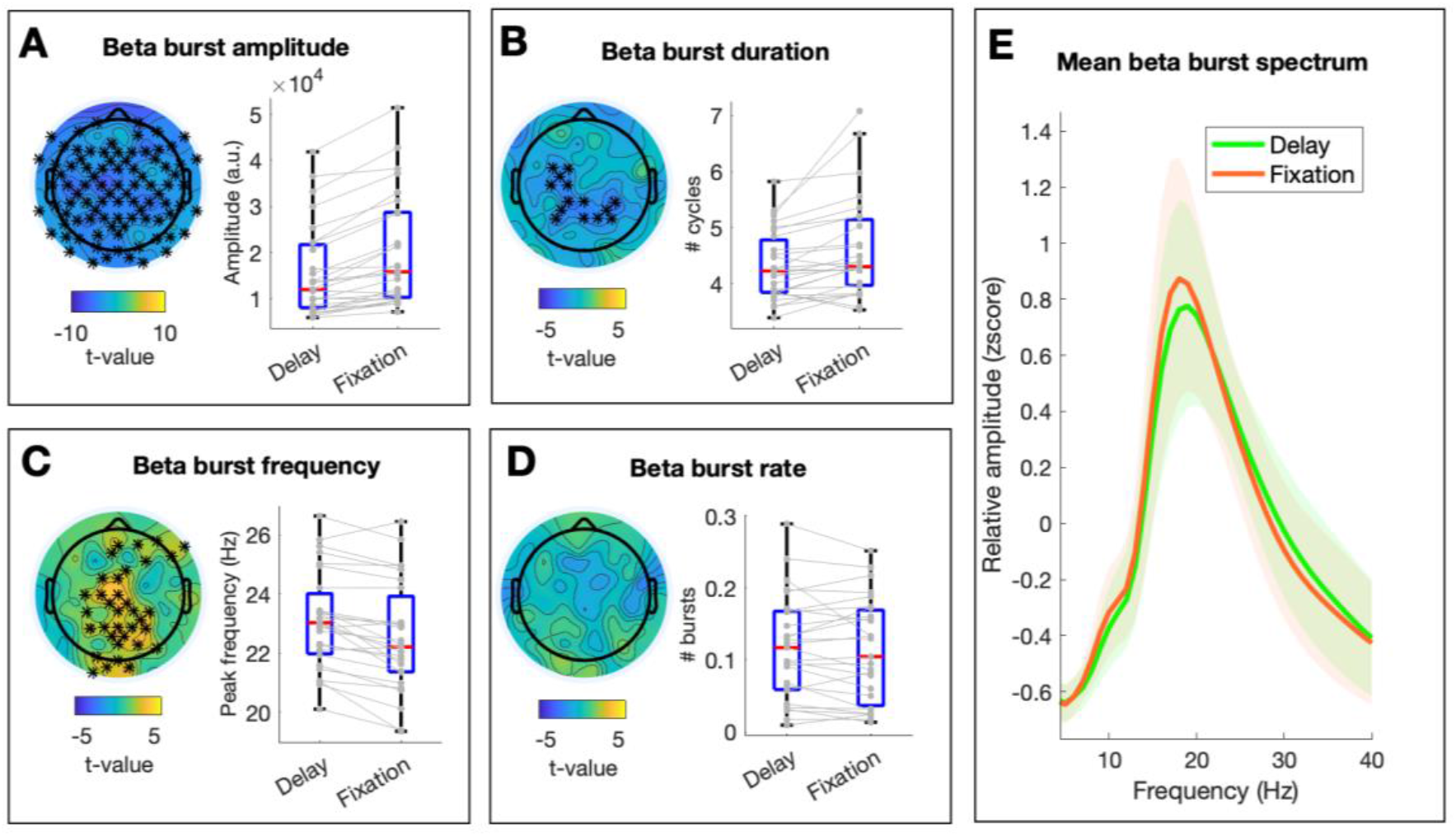
Changes in beta burst parameters during memory retention. **A-D)** The left panel depicts the topographical distribution of t-values for the comparison *Delay 1* vs *Fixation* for each beta burst parameter. Asterisks indicate significant clusters at *p*<0.025. The right panel shows individual values (in grey) and boxplots of significant clusters per condition. **E)** Mean frequency spectrum of detected beta bursts per condition.

### Beta burst modulations with cognitive load

Next, we contrasted beta burst parameters across load conditions. Cognitive load was associated with a significant decrease in beta burst amplitude (*t*_*cluster*_ = -176.49; *p*_*cluster*_ < 0.001; *d* = 0.32; **Figure 4A**), a significant decrease in beta burst duration (*t*_*cluster*_ = -248.39; *p*_*cluster*_ < .001; *d* = 0.67; **Figure 4B**), a significant increase in beta burst frequency (*t*_*cluster*_ = 165.20; *p*_*cluster*_ < .001; *d* = 0.36; **Figure 4C**) and a significant increase in beta burst rate (*t*_*cluster*_ = 87.56; *p*_*cluster*_ = .008; *d* = 0.19; **Figure 4D**). The topographical distribution of significant modulations was widespread for burst amplitude, duration and frequency, while effects on beta burst rates were more constrained to central electrodes. The average frequency spectra of beta bursts in each condition across electrodes are depicted in **Figure 4E**.

**Figure 4.**
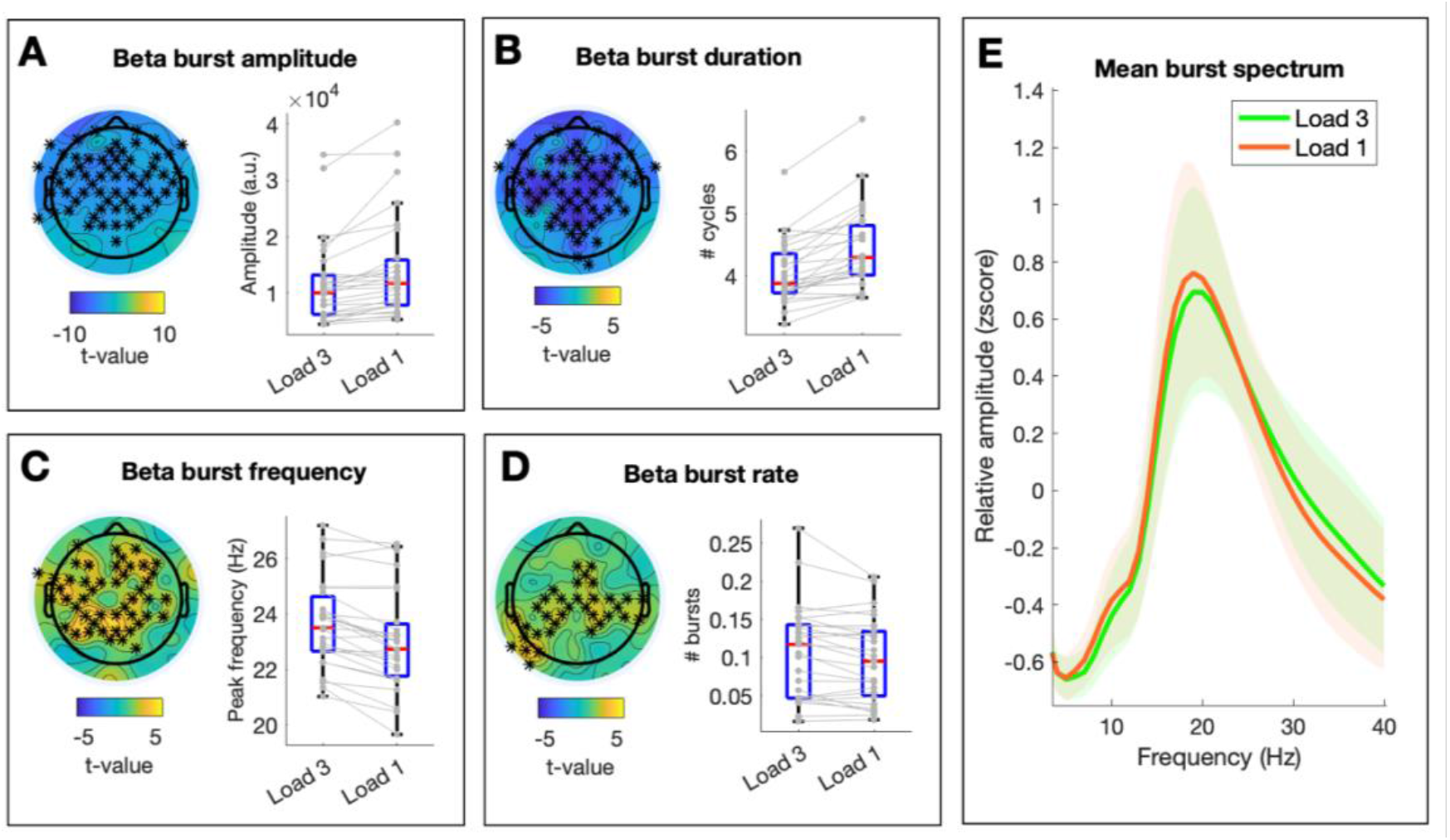
Changes in beta burst parameters associated with memory load. **A-D)** The left panel depicts the topographical distribution of t-values for the comparison *Load 3* vs *Load 1* for each beta burst parameter. Asterisks indicate significant clusters at *p*<0.025. The right panel shows individual values (in grey) and boxplots of significant clusters per condition. **E)** Mean frequency spectrum of detected beta bursts per condition.

### Beta burst modulations associated with memory manipulation

In order to assess the effect of memory manipulation on beta burst parameters, we contrasted switch and stay conditions during the second memory delay. Memory manipulation was associated with a significant decrease in beta burst amplitude (*t*_*cluster*_ = 75.41; *p*_*cluster*_ = .002; *d* = 0.18; **Figure 5A**), a significant decrease in beta burst duration (*t*_*cluster*_ = -103.00; *p*_*cluster*_ < .001; *d* = 0.40; **Figure 5B**), a significant increase in beta burst frequency (*t*_*cluster*_ = 29.55; *p*_*cluster*_ < .001; *d* = 0.30; **Figure 5C**) and a significant increase in beta burst rate (*t*_*cluster*_ = 52.30; *p*_*cluster*_ = .006; *d* = 0.11; **Figure 5D**). The topographical distribution of significant modulations was centro-parietal for burst amplitude, widespread for burst duration and limited to posterior electrodes for burst frequency and burst rate. The average frequency spectra of beta bursts in each condition across electrodes are depicted in **Figure 5E**.

**Figure 5.**
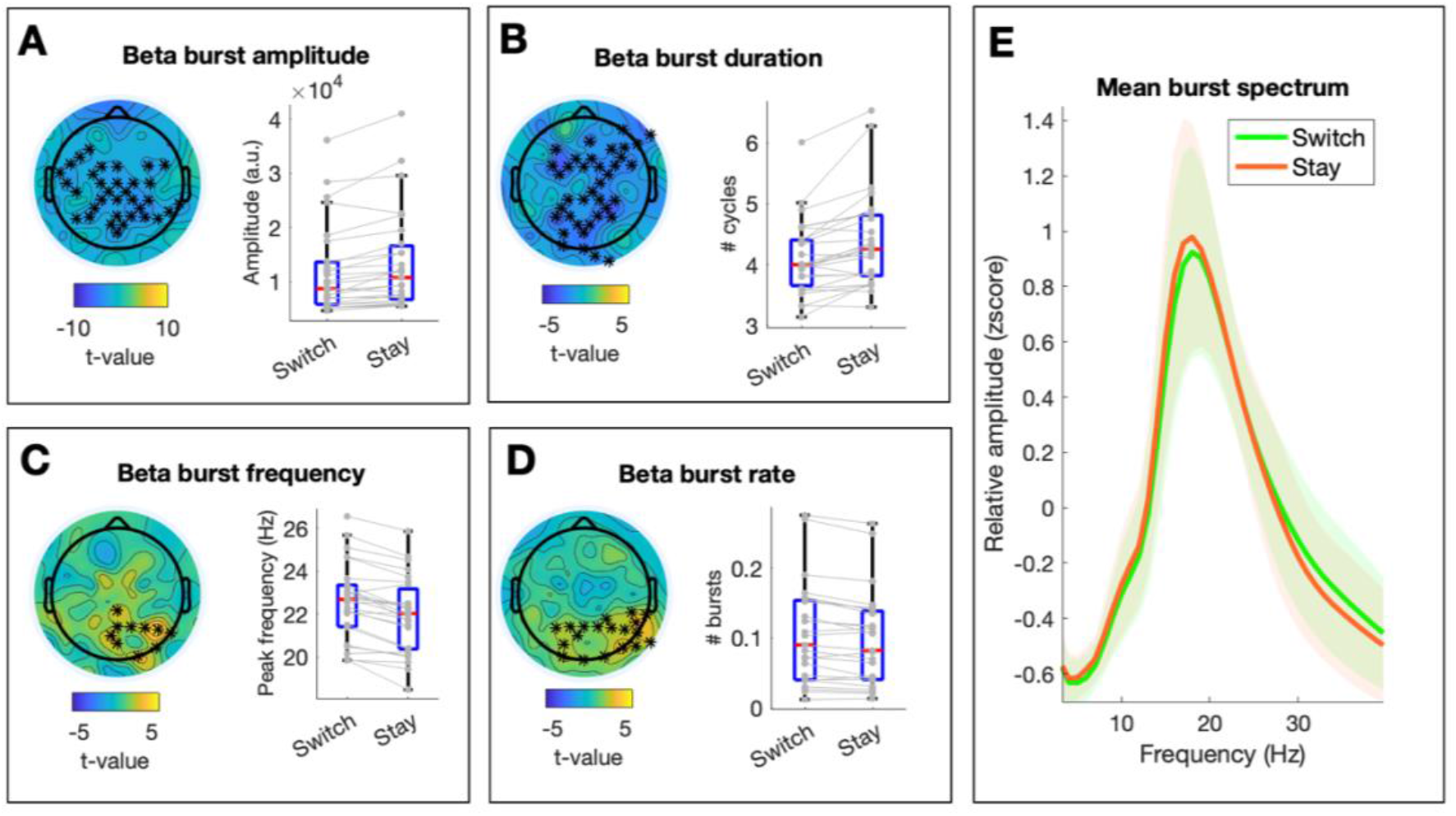
Changes in beta burst parameters associated with memory manipulation. **A-D)** The left panel depicts the topographical distribution of t-values for the comparison of conditions *Switch* and *Stay* for each beta burst parameter. Asterisks indicate significant clusters at *p*<0.025. The right panel shows individual values (in grey) and boxplots of significant clusters per condition. **E)** Mean frequency spectrum of detected beta bursts per condition.

### Relation between beta bursts and interindividual differences in performance

Finally, we asked whether beta burst parameters correlated with working-memory performance. No significant clusters were identified when assessing the relationship between accuracy and each of the four beta burst parameters during the first or second memory delay (all *p*_*cluster*_ > .05). Assessing interindividual differences in response times, we found a significant positive cluster for beta burst rate during the first delay (*t*_*cluster*_ = 34.90; *p*_*cluster*_ = .02; *mean r-value* = 0.47) (see **Figure 6**). This shows that participants with greater beta burst rates during the first memory delay tended to have slower response times. No significant clusters were identified for the remaining beta burst parameters for either delay (all *p*_*cluster*_ > .05).

**Figure 6.**
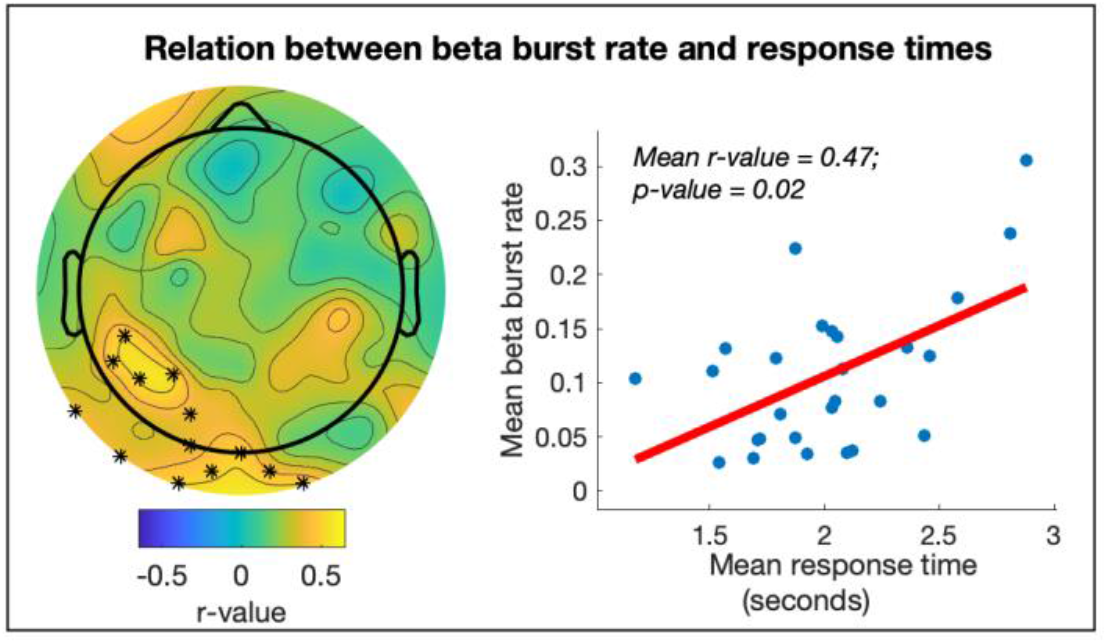
Relationship between beta burst rates and performance. The left panel depicts the topographical distribution of *r-values*. Asterisks indicate significant clusters at *p*<0.025. The right panel shows mean burst rates as a function of mean response times (each point depicts a participant).

## Discussion

Here, we investigated the functional relevance of beta oscillations in human working memory. We collected 96-electrode EEG data while participants performed a spatial working-memory task. Critically, we controlled for the possible influence of lower frequency rhythms with non-sinusoidal properties on beta-band dynamics. Specifically, we developed an algorithm that detects oscillatory bursts in the beta range that do not co-occur in time or space with more prominent rhythms in lower frequencies. Our results show significant modulations in several beta burst parameters during memory retention and manipulation. Both memory load and memory manipulation were associated with decreased beta burst amplitude, decreased beta burst duration, increased beta burst rate and increased beta burst peak frequency. In addition, we found that only beta burst rate showed a significant relation with performance: participants with slower response times tended to have a higher rate of beta bursts during the memory delay. Together, these results show that beta oscillations that cannot be attributed to non-sinusoidal properties of lower frequency rhythms are significantly modulated during working memory and predict performance.

Previous literature on the role of beta oscillations in human working memory focused almost exclusively on amplitude modulations, with seemingly contradictory results (Pavlov & Kotchoubey, 2020b). Memory load and memory manipulation have been associated with both increases (Chen & Huang, 2016; Deiber et al., 2007; Tallon-Baudry et al., 1998) and decreases in beta amplitude (Erickson et al., 2019; Nasrawi & van Ede, 2022; Pavlov & Kotchoubey, 2020a; Proskovec et al., 2018). There are two main difficulties that make these previous results hard to interpret. First, given the burst-like nature of beta oscillations (Jones, 2016; van Ede et al., 2018), changes in amplitude as estimated with conventional analyses could be confounded by changes in other parameters such as rate or duration of beta bursts (Donoghue et al., 2022). Secondly, putative modulations in beta amplitude could also be produced artifactually due to changes in a lower frequency rhythm with non-sinusoidal properties (Cole & Voytek, 2017; Schaworonkow & Nikulin, 2019). Here, we control for the influence of these two factors when estimating beta amplitude, and show that beta decreases with cognitive load and during memory manipulation. Thus, our beta burst algorithm might help reconcile previous results if applied to other data sets (see Methods for access to the analysis code). By controlling for the possible influence of lower frequency rhythms and separating the contribution of different beta burst parameters, we can robustly assess whether previous inconsistencies in the literature are due to the analytical approach and/or to other factors such as the modality or difficulty of the adopted working-memory task.

The here reported modulations in the amplitude and frequency of beta oscillations can be interpreted as changes in cortical excitability. On the one hand, the amplitude of oscillations in the alpha/beta range in humans has been associated with decreased broadband high frequency activity (BHA; Iemi et al., 2022), which is thought to provide a measure of local neuronal excitability (Leszczyński et al., 2020; Ray & Maunsell, 2011). On the other hand, modelling work suggests that increases in the peak frequency of neural oscillations are accompanied by increases in spiking activity of individual neurons (Mierau et al., 2017). In light of these studies, a decrease in beta amplitude and increase in beta frequency with memory load and during memory manipulation might be interpreted as a general increase in the excitability of task-relevant cortical areas. The topographical differences between load and manipulation effects (i.e., manipulation effects are more posterior; see **Figures 4–5**) could then be due to the recruitment of different areas for these two cognitive operations (Jablonska et al., 2020; Veltman et al., 2003).

Based on our results and previous theoretical accounts (Jensen & Mazaheri, 2010; Klimesch et al., 2007; Spitzer & Haegens, 2017), we speculate that “sustained” (i.e., long duration and low rate) and “transient” (i.e., short duration and high rate) oscillatory activity might reflect different neural processes. We propose that while sustained oscillations reflect functional inhibition, transient bursts are a cause (or consequence; see Schneider et al., 2021) of a set of task-relevant neural populations being (re)activated (Spitzer & Haegens, 2017), with the specific frequency of oscillatory activity related to the size of the neural populations being “inhibited” or “reactivated” (i.e., lower frequencies for bigger networks; see von Stein & Sarnthein, 2000). Theta oscillations are an intuitive example of this dual role of neural oscillations in humans, as they appear in the transition to sleep in a more sustained manner (putatively reflecting cortical inhibition; Canales-Johnson et al., 2020; Strijkstra, Beersma, Drayer, Halbesma, & Daan, 2003) and transiently during working-memory tasks in prefrontal areas (which are known to be task-relevant; see Müller & Knight, 2006). This tentative view might reconcile the seeming discrepancy between reports of beta dynamics that suggest an “inhibitory” role, and those that seem to reflect the formation of neural ensembles (see Spitzer & Haegens, 2017). Together, we hypothesize that the here reported decreases in duration and increases in rate of beta bursts during working-memory retention and manipulation are reflective of the transient reactivation of content-specific neural circuits. Although there is some evidence for this function of beta bursts from recordings in monkeys (Buschman et al., 2012; Rassi et al., 2023), the analysis of human intracranial EEG data that includes the simultaneous recording of both local field potentials and spiking activity is necessary to fully assess our predictions.

From the four extracted beta burst parameters, only rate (i.e., the mean number of bursts) was significantly associated with interindividual differences in performance. Specifically, we found that participants with higher beta burst rates during the first memory delay tended to have slower response times, which is in line with a recent report (Rassi et al., 2022). These results should be interpreted with caution, as response times in working memory tasks depend on a wide variety of factors such as cognitive strategy, level of arousal and working memory capacity (Miyake, 2001; Robison & Brewer, 2020). Since beta burst rates also increased with task difficulty (i.e., load and manipulation effects), we speculate that the task was easier for faster participants, which might be why they showed a relatively lower number of beta bursts. A bigger sample size with greater interindividual variability in performance and a thorough assessment of trait and state-like interindividual differences (e.g., a combination of questionnaires and experience sampling; Mrazek et al., 2013) is needed to robustly assess the relationship between beta bursts and interindividual differences in working-memory performance.

Finally, it is important to underline that several algorithms to detect oscillatory bursts have been proposed (Bonaiuto et al., 2021; Neymotin et al., 2022; Seymour et al., 2022; Shin et al., 2017; Whitten et al., 2011), and that each of them offers specific advantages. An important factor for the precise detection of oscillatory bursts is the definition of an amplitude threshold. In line with a recently developed algorithm (Szul et al., 2022), we here use the estimate of aperiodic activity as a power threshold, instead of a global threshold of beta amplitude (Bonaiuto et al., 2021; Sherman et al., 2016; Shin et al., 2017). This strategy was adopted to also detect low-amplitude oscillatory bursts, which might be of functional relevance (Schürmann & Başar, 2001). In this regard, note that the algorithm developed here specifically aimed to control for the influence of non-sinusoidal low-frequency rhythms on beta oscillations (Schaworonkow, 2023; Schaworonkow & Nikulin, 2019), because, to our knowledge, this factor was not taken into account in previously proposed algorithms. However, other algorithms offer other features, such as the estimation of waveform shape or the detection of rhythms that change in instantaneous frequency (Neymotin et al., 2022; Szul et al., 2022), which might make them more adequate for other research questions.

